# Reconstruction of Small Subunit Ribosomal RNA from High-Throughput Sequencing Data: A Comparative Study of Metagenomics and Total RNA Sequencing

**DOI:** 10.1101/2022.08.26.505493

**Authors:** Christopher A. Hempel, Shea E. E. Carson, Tyler A. Elliott, Sarah J. Adamowicz, Dirk Steinke

## Abstract

The small subunit (SSU) ribosomal RNA (rRNA) is the most commonly used marker for the identification of microbial taxa, but its full-length reconstruction from high-throughput sequencing (HTS) data remains challenging, especially for complex and diverse environmental samples. Metagenomics and total RNA sequencing (total RNA-Seq) are target-PCR-free HTS methods that are used to characterize microbial communities and simultaneously reconstruct SSU rRNA sequences. However, more testing is required to determine and improve their effectiveness. In this study, we processed metagenomics and total RNA-Seq data retrieved from a commercially available mock microbial community using 112 combinations of commonly used data-processing tools, determined SSU rRNA reconstruction completeness of both sequencing methods for each species in the mock community, and analyzed the impact of data-processing tools on SSU rRNA and genome completeness. Total RNA-Seq allowed for the complete or near-complete reconstruction of all mock community SSU rRNA sequences and outperformed metagenomics. SSU rRNA completeness of metagenomics strongly correlated with the genome size of mock community species. The impact of data-processing tools was overall low, although certain tools resulted in significantly lower SSU rRNA completeness. These results are promising for the high-throughput reconstruction of novel full-length SSU rRNA sequences and could advance the simultaneous application of multiple -omics approaches in routine environmental assessments to allow for more holistic assessments of ecosystems.

## 1 Introduction

Microbial organisms (prokaryotes and unicellular eukaryotes) make up most of the biodiversity on our planet, with the global number of microbial species estimated at up to one trillion (Locey & Lennon, 2016). These organisms play crucial roles in every biome on Earth, including the microbiomes within other organisms (e.g., humans), and they are very sensitive to environmental change. In fact, the microbial community composition of a particular environment can provide us with important information about the state and health of the environment (Cordier, Lanzén, Apothéloz-Perret-Gentil, Stoeck, & Pawlowski, 2019; Pawlowski, Lejzerowicz, Apotheloz-Perret-Gentil, Visco, & Esling, 2016; Proctor et al., 2019; Sagova-Mareckova et al., 2021; Smith et al., 2015).

Determining this community composition requires the identification of its member taxa. Since most microbes lack diagnostic traits and are too abundant and diverse to be identified morphologically, their identity is typically determined by DNA-based methods (Pawlowski et al., 2012; Woese, 1987), which require DNA reference databases. Ideally, such databases contain full-length genomes of all microbial taxa on Earth to allow for taxonomic annotations at the highest possible resolution. However, in reality, most microbial taxa are unknown, and even for known taxa, the number of available full-length or even partial genomes is very low, since the high-throughput reconstruction of full-length genomes has only recently become possible. As a consequence, the largest and most widely used curated database for genomes, NCBI RefSeq, only comprises genomes of 71,311 microbial organisms (release 213: https://ftp.ncbi.nlm.nih.gov/refseq/release/release-notes/RefSeq-release213.txt). The largest project cataloging all microbes on Earth (Earth Microbiome Project: http://www.earthmicrobiome.org) has led to the reconstruction of 52,515 microbial genomes to date (Nayfach et al., 2021). This clearly shows that the reconstruction of all microbial genomes or at least a larger segment of microbial diversity will require much more time.

For the time being, a more feasible and common approach to identifying microbial taxa is to focus on specific genes. The small subunit (SSU) ribosomal RNA (rRNA) gene (16S rRNA for prokaryotes and 18S rRNA for eukaryotes) is the primary marker gene for microbes. SSU rRNA reference databases are also far from complete, but the most commonly used database, SILVA, contains 510,508 full-length, non-redundant SSU rRNA gene sequences (release 138.1 SSU Ref NR 99: https://www.arb-silva.de/documentation/release-1381/), and the number is growing quickly. Full-length SSU rRNA reference sequences are paramount since shorter sequences, which are generated by short-read amplicon-sequencing, can lead to inaccurate taxonomic identifications (Johnson et al., 2019; Yarza et al., 2014). The traditional approach to reconstructing full-length SSU rRNA reference sequences involves cloning and Sanger sequencing, which is costly and comes with limited throughput. Advancements in sequencing technology gave rise to new approaches at lower costs and with higher throughput. Such high-throughput sequencing (HTS) approaches simultaneously allow us to characterize the microbial community composition of environmental samples and are therefore extremely powerful to advance our understanding of microbial communities.

Multiple HTS approaches have been developed to date, and they can primarily be categorized into short-read sequencing (including synthetic long-read sequencing) and true long-read sequencing. True long-read sequencing reads have an average accuracy of >99.5% (Pacific Biosciences) or 92–94% (Oxford Nanopore Technologies) at an average length of 10–25 kb (Delahaye & Nicolas, 2021; Hon et al., 2020), which is multiple times the length of the complete SSU rRNA (approximately 1.46 kb on average, based on the average length of all sequences in SILVA release 138.1 SSU Ref NR 99). However, although true long-read sequencing has been applied successfully for some microbial community analysis (Singer et al., 2016; Tedersoo, Albertsen, & Anslan, 2021), it is limited by the available read depth of long-read instruments. This makes it less scalable and currently not applicable for the study of complex microbial communities in comparison with short-read or synthetic long-read sequencing.

Metagenomics and metatranscriptomics are short-read sequencing methods without target PCR that are generally applied to analyze the presence and expression of functional genes within communities (Almeida & De Martinis, 2019; Bashiardes, Zilberman-Schapira, & Elinav, 2016; Shakya, Lo, & Chain, 2019; Wooley, Godzik, & Friedberg, 2010). Both involve random fragmentation of genomes and transcripts into shorter sequences, thereby randomly covering smaller fragments of the SSU rRNA of organisms in a sample. These fragments can be utilized for SSU rRNA sequence reconstruction and taxonomic identification of communities simultaneously.

Metagenomics targets all DNA in a sample and therefore has broad genomic coverage, which is important for functional analyses of communities. However, this also means that the coverage of particular genes of interest, such as the SSU rRNA gene, is low. Consequently, SSU (and LSU) rRNA sequences can make up as little as 0.05–1.4% of metagenomics sequences (Logares et al., 2014; Yilmaz et al., 2011). Targeted tools have been developed to extract and reconstruct SSU rRNA sequences from metagenomics datasets (Bengtsson-Palme et al., 2015; Fan, McElroy, & Thomas, 2012; Miller, Baker, Thomas, Singer, & Banfield, 2011; Pericard, Dufresne, Couderc, Blanquart, & Touzet, 2018; Yuan, Lei, Cole, & Sun, 2015; Zeng, Wang, Wang, Zhou, & Chen, 2017), but overall, the low proportion of SSU rRNA makes the approach inefficient for SSU rRNA reconstruction and taxonomic community analysis.

An alternative approach is total RNA sequencing (total RNA-Seq) (Bang-Andreasen et al., 2020; F. Li & Guan, 2017; F. Li et al., 2016), also termed double-RNA approach (Urich et al., 2008), metatranscriptomics analysis of total rRNA (Turner et al., 2013), total RNA metatranscriptomics (Xue, Lanzén, & Jonassen, 2020), or total RNA-seq-based metatranscriptomics (F. Li & Guan, 2017), all referring to metatranscriptomics without an mRNA enrichment step. As 80–98% of cellular RNA consists of rRNA (Peano et al., 2013; Westermann, Gorski, & Vogel, 2012), SSU (and LSU) rRNA can make up 37–71% of all total RNA sequences (Elekwachi, Wang, Wu, Rabee, & Forster, 2017; Yu & Zhang, 2012). This means that a large portion of total RNA-Seq data can be used for full-length assembly of SSU rRNA.

Several studies took total RNA-Seq a step further and combined it with synthetic long-read sequencing (Dueholm et al., 2020; Karst et al., 2018; Tedersoo et al., 2021). Synthetic long-read sequencing is a modification of short-read sequencing in which each molecule is tagged with unique molecular identifiers (UMIs), amplified, randomly split, and sequenced with conventional short-read sequencing technology. Fragments of the same molecule are linked through the tags, and complete molecules can be reconstructed from linked short sequences through *de novo* assembly, hence the term synthetic long reads. Karst et al. (2018) size-selected total RNA for SSU rRNA and applied synthetic long-read sequencing to generate over a million SSU rRNA sequences, of which almost 45,000 were complete. Although this approach is effective, it adds additional costs, time, and room for bias. Due to the growing interest in applying multiple -omics approaches to environmental samples to generate a more holistic picture of ecosystems (Cordier et al., 2021, 2019; Leese et al., 2018; Uyaguari-Diaz et al., 2016), less specialized methods might be easier to implement, especially for routine application. Total RNA-Seq without size selection and UMI tagging follows the same protocol as conventional metatranscriptomics after mRNA enrichment, and both methods could be implemented simultaneously without additional modifications.

A few studies compared the performance of total RNA-Seq and metabarcoding (Lanzén et al., 2011; Yan et al., 2018) or metagenomics (Hempel et al., 2022; Lanzén et al., 2011; Shi, Tyson, Eppley, & Delong, 2011; Urich et al., 2014; Uyaguari-Diaz et al., 2016) for the analysis of microbial community composition. However, the use of total RNA-Seq to analyze microbial communities remains rare, and a comparison of total RNA-Seq and metagenomics in terms of SSU rRNA reconstruction is lacking, although the results of such a comparison could impact our ability to categorize global biodiversity and analyze microbial communities.

In an earlier study, we were able to show that total RNA-Seq characterized a mock microbial community consisting of 10 species more accurately than metagenomics at almost one order of magnitude lower sequencing depth (Hempel et al., 2022). For the present study, we use the same data to test the performance of metagenomics and total RNA-Seq in terms of SSU rRNA completeness, i.e., the portion of the SSU rRNA that can be successfully reconstructed. Additionally, we determine and compare the genome completeness of both approaches to highlight the advantages and disadvantages of either approach. Lastly, since total RNA-Seq has not yet been extensively tested, we also determine the impact of commonly used data-processing tools on SSU rRNA reconstruction, as it has been repeatedly shown that results based on HTS are heavily influenced by the choice of bioinformatics tools (Bashiardes et al., 2016; Knight et al., 2018; McIntyre et al., 2017; Quince, Walker, Simpson, Loman, & Segata, 2017; Shakya et al., 2019; Vollmers, Wiegand, & Kaster, 2017).

## 2 Methods

The overall study design is shown in Figure 1, and further details are given in the balance of this section.

**Figure 1:**
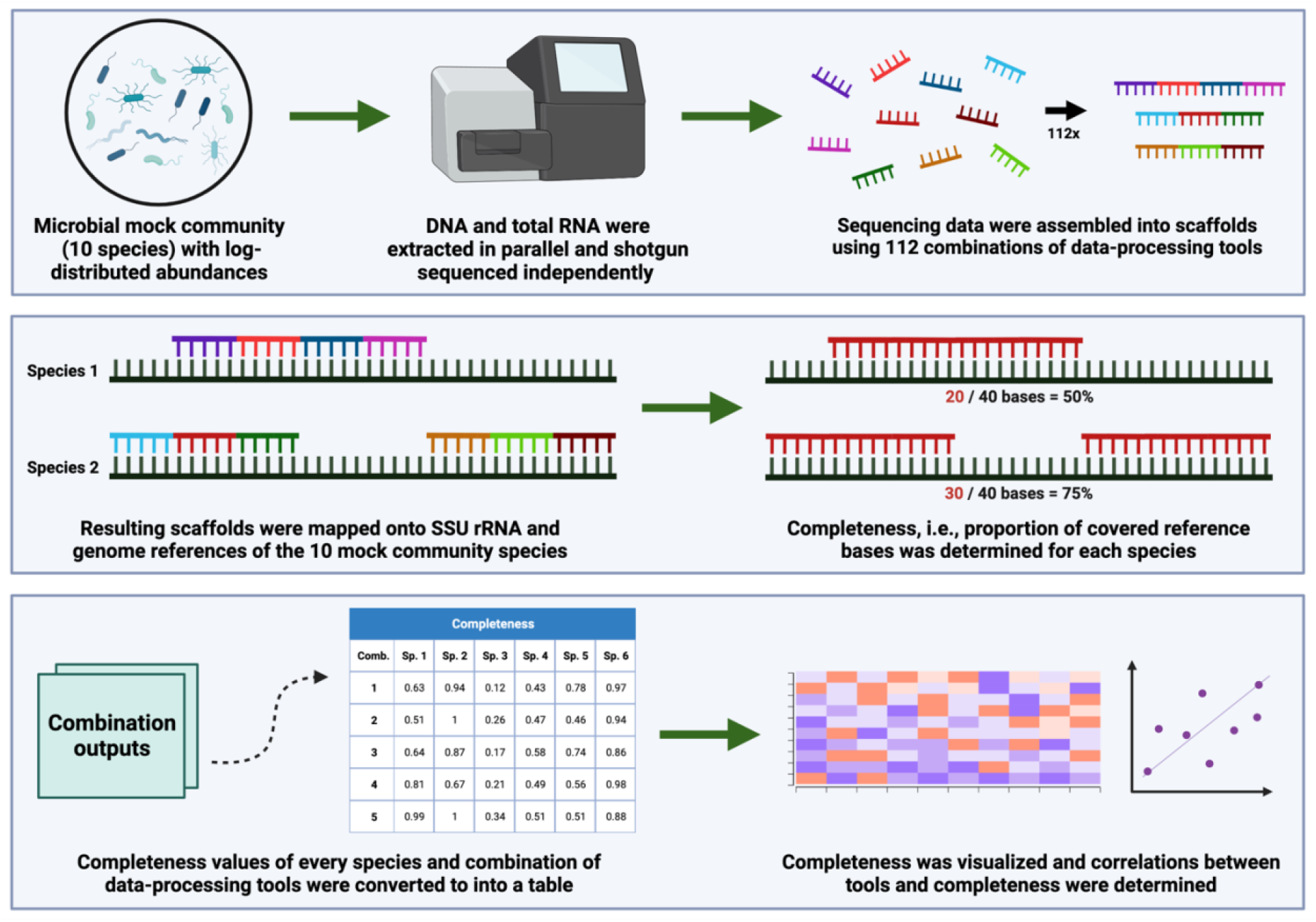
Summary of the study design. The DNA and total RNA of a mock microbial community were extracted and shotgun-sequenced on an Illumina MiSeq, representing two sequencing methods (metagenomics and total RNA-Seq). Sequencing data were assembled into scaffolds using 112 combinations of common data-processing tools. Scaffolds were mapped onto SSU rRNA and genome references of the 10 mock community species, and the completeness, i.e., proportion of covered bases was determined for each species. The completeness values for all combinations of data-processing tool were converted into a table and used for visualizations and statistical analyses. This figure was created with BioRender.com.

### 2.1 Mock microbial community

We used a commercially available mock microbial community (ZymoBIOMICS Microbial Community Standard II (Log Distribution); Zymo Research; Irvine; CA U.S.A.), consisting of eight bacterial species (three gram-negative and five gram-positive) and two yeast species with log-distributed species abundances based on genomic DNA quantity (Figure 2). The mock community was preserved in DNA/RNA Shield (Zymo Research; Irvine; CA U.S.A.) by the manufacturer to inactivate cells while preserving DNA and RNA. We generated three simulated water sample replicates by adding 130 μl of the mock community containing approximately 381 ng of total DNA to 50 mL ultrapure water respectively.

**Figure 2:**
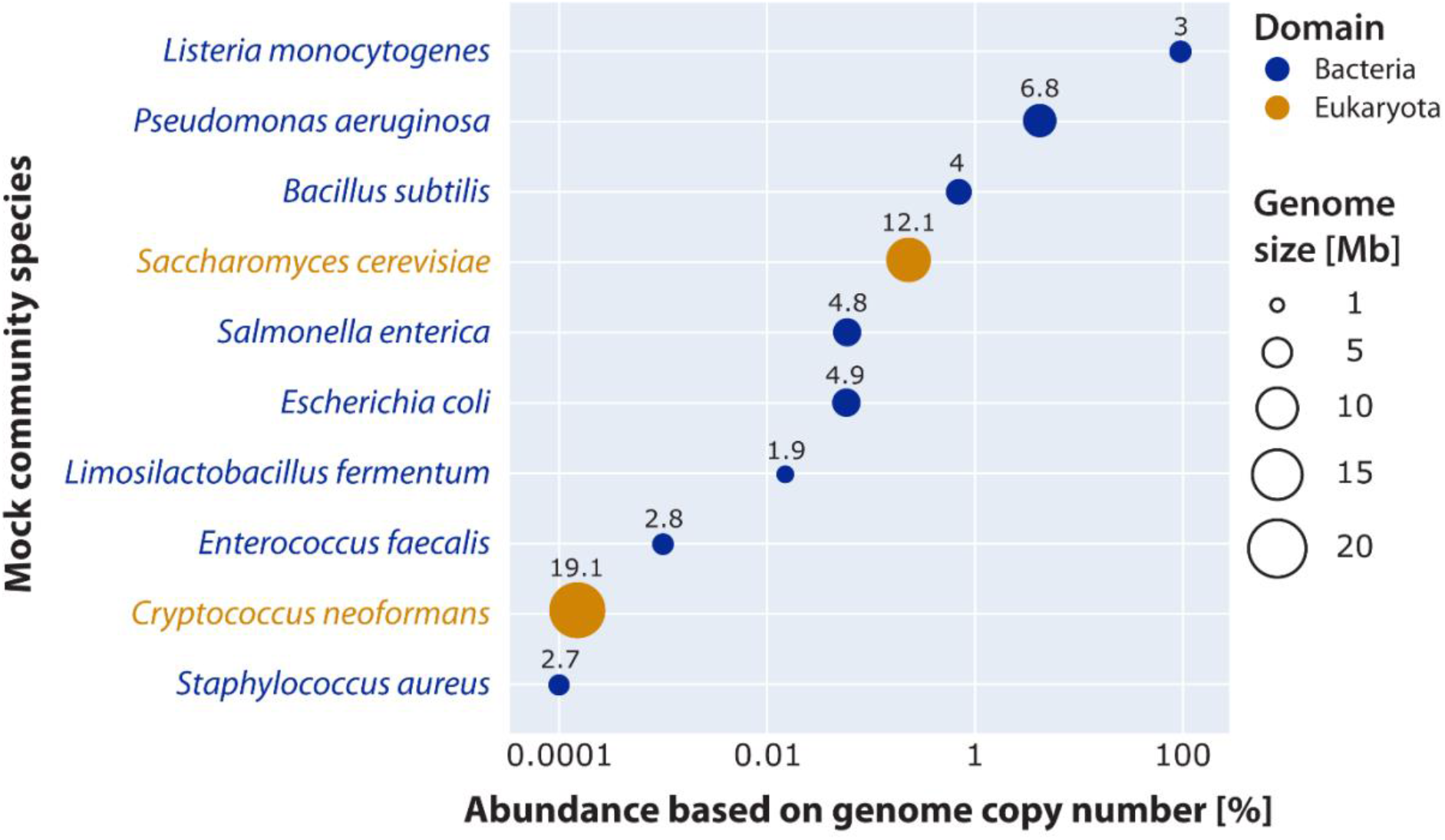
Genome size and relative abundance of mock community species. Relative abundances are based on genome copy numbers, as recommended by the manufacturer for studies that involve shotgun sequencing. The size of and numbers above bubbles indicate genome sizes in megabases (Mb).

### 2.2 Laboratory and bioinformatics processing

The data used in this study originate from an earlier study, in which we applied total RNA-Seq and metagenomics and investigated 672 combinations of data-processing tools to identify the best-performing sequencing method and combination of tools to characterize taxonomically a mock microbial community (Hempel et al., 2022). Details of the laboratory and bioinformatics processing steps can be found there (Hempel et al., 2022). In summary, we co-extracted DNA and RNA in parallel using a modified version of the Quick-DNA/RNA Microprep Plus Kit (Zymo Research; Irvine; CA U.S.A.). DNA and RNA library prep and sequencing were performed by a service provider (Génome Québec; Montreal; QC Canada), and mRNA enrichment was skipped prior to RNA library prep to sequence total RNA. During library prep, normalization was performed by processing equal volumes of samples instead of equal concentrations of samples so that the relative number of reads per sample mirrored the relative amount of DNA/RNA, avoiding an over-or underrepresentation of samples with higher or lower amounts of DNA/RNA. The libraries were sequenced on an Illumina MiSeq together with samples from another project (2,428,038 paired-end reads in total; Bioproject number: PRJNA819997; SRA accession number: SRR18488964–SRR18488973). Metagenomics replicates received a much higher number of reads than total RNA-Seq replicates, and to allow for appropriate comparisons between both approaches, we subsampled the reads of metagenomics replicates to match the number of reads of total RNA-Seq replicates. As DNA and RNA were coextracted, we subsampled each metagenomics replicate to the number of reads of the corresponding total RNA-Seq replicate. Random subsampling was performed ten times, and the subsamples were processed independently.

Sequence processing for this study was divided into three data-processing steps: trimming and quality filtering, rRNA sorting, and assembly. For trimming and quality filtering, four PHRED score cut-offs were applied (PHRED ≤5, ≤10, ≤15, and ≤20) to trim the end of reads using Trimmomatic v0.39 (Bolger, Lohse, & Usadel, 2014). For rRNA sorting, three approaches were applied to sort reads into rRNA and non-rRNA reads: alignment-based with SortMeRNA v4.0.0 (Kopylova, Noé, & Touzet, 2012), Hidden Markov model-based with barrnap v0.9 (Seemann, unpublished, https://github.com/tseemann/barrnap, accessed on 18 Jun 2021), and kmer-based with rRNAFilter v1.1 (Wang, Hu, & Li, 2017). For each approach, non-rRNA reads were subsequently excluded. Additionally, we omitted rRNA sorting and used all reads instead, leading to four rRNA sorting approaches in total. For assembly, we tested seven assemblers for both metagenomics and total RNA-Seq reads: SPAdes, metaSPAdes v3.14.1 (Nurk, Meleshko, Korobeynikov, & Pevzner, 2017), MEGAHIT v1.2.9 (D. Li, Liu, Luo, Sadakane, & Lam, 2015), IDBA-UD v1.1.1 (Peng, Leung, Yiu, & Chin, 2012), Trinity v2.10.0 (Grabherr et al., 2013), rnaSPAdes v3.14.1 (Bushmanova, Antipov, Lapidus, & Prjibelski, 2019), IDBA-tran v1.1.1 (Peng et al., 2013), and Trans-ABySS v2.0.1 (Robertson et al., 2010). Trinity (Grabherr et al., 2013) is another widely used, metatranscriptomics-optimized assembler, but despite thorough efforts to run Trinity with adjusted RAM settings as recommended by the developers (https://trinityrnaseq.github.io/performance/mem.html), we ultimately failed to operate it consistently. Furthermore, there are multiple assemblers specifically designed for SSU rRNA assembly from metagenomics and even total RNA-Seq data, such as EMIRGE (Miller et al., 2011), MATAM (Pericard et al., 2018), MetaRib (Xue et al., 2020), RAMBL (Zeng et al., 2017), and REAGO (Yuan et al., 2015), but although we tried to run the two most recent of them, MATAM and MetaRib, MATAM runs consistently failed and we were not able to successfully install MetaRib (more specifically EMIRGE, which is a requirement for MetaRib). Therefore, we could not include Trinity or SSU-rRNA-optimized assemblers into our benchmarking, and the lack of user friendliness or flexibility of these assemblers should be taken into account in future studies.

Combining all data-processing tools in the three steps resulted in 112 combinations of tools that were applied to both metagenomics and total RNA-Seq data, respectively (224 in total). The code to run all combinations is available on GitHub (https://github.com/hempelc/metagenomics-vs-totalRNASeq-reference-comparison).

### 2.3 Determining SSU rRNA and genome completeness of mock community species

For each mock community species, we determined both SSU rRNA and genome completeness of the scaffolds generated in each of the 224 combinations separately. To determine completeness, references were required for each mock community species. Zymo Research provides full-length SSU rRNA and genome references for all mock community species, but SSU rRNA references of some species include multiple sequences to cover strain variants, and the genome references for the two eukaryotic species (*Saccharomyces cerevisiae* and *Cryptococcus neoformans*) consisted of draft instead of curated genomes. Therefore, we aligned all SSU rRNA references of species with multiple reference sequences, respectively, using Geneious Prime v2022.1.1 (https://www.geneious.com) and the mafft plugin v7.450 (Katoh & Standley, 2013) with default parameters, extracted the consensus sequence at a 100% threshold to include all ambiguities, and used the consensus sequences as SSU rRNA references. We also downloaded the reference genomes of the eukaryotic species from RefSeq (GCF_000146045.2 for *S. cerevisiae* and GCF_000091045.1 for *C. neoformans*) and used those as reference genomes instead of the draft genomes provided by Zymo Research.

Both SSU rRNA and genome references for the ten mock microbial community species were indexed for mapping using the *index* function of BWA v0.7.17 (H. Li & Durbin, 2009) with default parameters. The scaffolds resulting from each combination of tools were mapped onto both the indexed SSU rRNA and the genome reference of each species using the BWA-MEM algorithm of BWA with default parameters. We determined the completeness of each reference using the *coverage* function of samtools v1.10 (H. Li et al., 2009). Specifically, as the eukaryotic reference genomes consist of multiple chromosomes, we determined the total number of covered bases and divided it by the total number of bases to generate the relative completeness of each reference. This was done separately for each replicate, species, SSU rRNA and genome reference, and combination of tools. Since metagenomics replicates were randomly subsampled ten times, we determined the completeness of the metagenomics replicates by taking the mean average across the ten subsamples. We summarized the completeness across metagenomics and total RNA-Seq replicates, respectively, by taking the mean average across the three replicates. The references used and the code to determine completeness are available on GitHub (https://github.com/hempelc/metagenomics-vs-totalRNASeq-reference-comparison).

### 2.4 Statistical analysis and visualization

All further data processing, statistical analysis, and visualization were performed using Python v3.7.9 (Van Rossum & Drake, 2009). The full code is available on GitHub (https://github.com/hempelc/metagenomics-vs-totalRNASeq-reference-comparison) and involves the Python modules Pandas v1.3.5 (McKinney, 2010), NumPy v1.21.3 (Harris et al., 2020), and Plotly v5.6.0 (Plotly Technologies Inc., 2015).

We sorted SSU rRNA and genome completeness across species by data-processing tools and visualized differences in completeness using heatmaps. We visually observed a correlation between genome size and metagenomics SSU rRNA completeness, so we calculated the Pearson correlation coefficient between genome size and mean average SSU rRNA completeness across data-processing tools for both metagenomics and total RNA-Seq to test for statistically significant correlations using the pearsonr function of the Python module SciPy v1.7.1 (Virtanen et al., 2020).

Additionally, we calculated the Pearson correlation coefficient between abundance and mean average SSU rRNA completeness to test if species abundance correlated with SSU rRNA completeness.

As the genome completeness of all but one species was close to 0%, we only determined the impact of data-processing tools on SSU rRNA completeness. For each species, we performed a one-way ANOVA on SSU rRNA completeness obtained through all applied PHRED score cut-offs, rRNA sorting methods, or assembly tools, respectively, to determine if methods applied within each data-processing step significantly differed from another in terms of SSU rRNA completeness using the f_oneway function of SciPy. For some species, rRNA sorting methods and assembly tools differed significantly from another in terms of SSU rRNA completeness, which is why we converted them into binary dummy variables and tested for significant correlations between each tool and SSU rRNA completeness by calculating the Point-biserial correlation coefficient, which measures the strength of the association between continuous and binary variables, using the pointbiserialr function of SciPy.

## 3 Results

We found differences between metagenomics and total RNA-Seq when determining whether the methods within each data-processing step differed statistically from each other in terms of SSU rRNA completeness (Figure 3). The choice of methods for trimming and quality filtering did not significantly impact completeness for metagenomics or total RNA-Seq. However, for metagenomic data, the utilized rRNA sorting tools significantly impacted completeness across all species (p < 0.01). In contrast, for total RNA-Seq, rRNA sorting tools had no effect on SSU rRNA completeness for all species (p ≥ 0.94) except for the two eukaryotes (*S. cerevisiae:* p = 0.14, *C. neoformans:* p = 0.03). The assembly tools had a significant impact on SSU rRNA completeness (p < 0.01) across all species for total RNA-Seq and most species for metagenomics.

**Figure 3:**
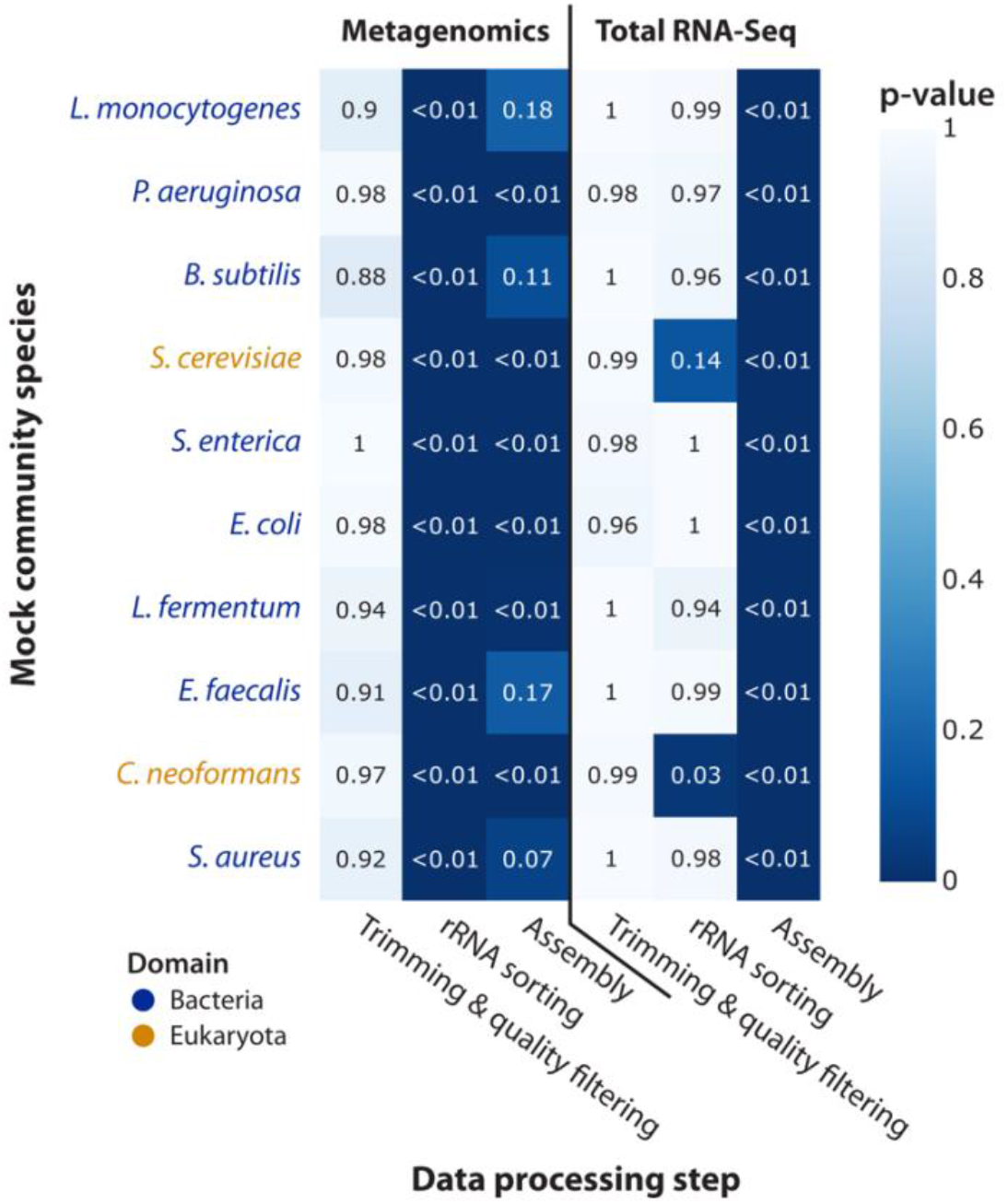
Impact of data-processing steps on SSU rRNA completeness. The x-axis represents the data-processing steps, the y-axis represents the ten mock community species, and the numbers and colour bar show the p-value of one-way ANOVAs between the tools of each step. Dark colours indicate that a particular processing step had a strong impact upon SSU rRNA completeness.

In terms of SSU rRNA completeness, the results varied considerably between the two sequencing methods (Figure 4). For metagenomics, SSU rRNA completeness was very low for both eukaryotic species, *S. cerevisiae* and *C. neoformans*, and low for a few bacterial species independent from the data-processing tools used. Statistically, SSU rRNA completeness was significantly and strongly negatively correlated to genome size and not correlated to species abundance. In contrast, for total RNA-Seq, SSU rRNA sequences were near-complete for all but three species across most data-processing tools, with some exceptions for the assemblers metaSPAdes and IDBA-tran. Statistically, SSU rRNA completeness was not correlated with genome size or species abundance for total RNA-Seq, although three of the four species with the lowest abundance were the only species without near-complete SSU rRNA sequences (*L. fermentum*, *E. faecalis*, and *C. neoformans*).

**Figure 4:**
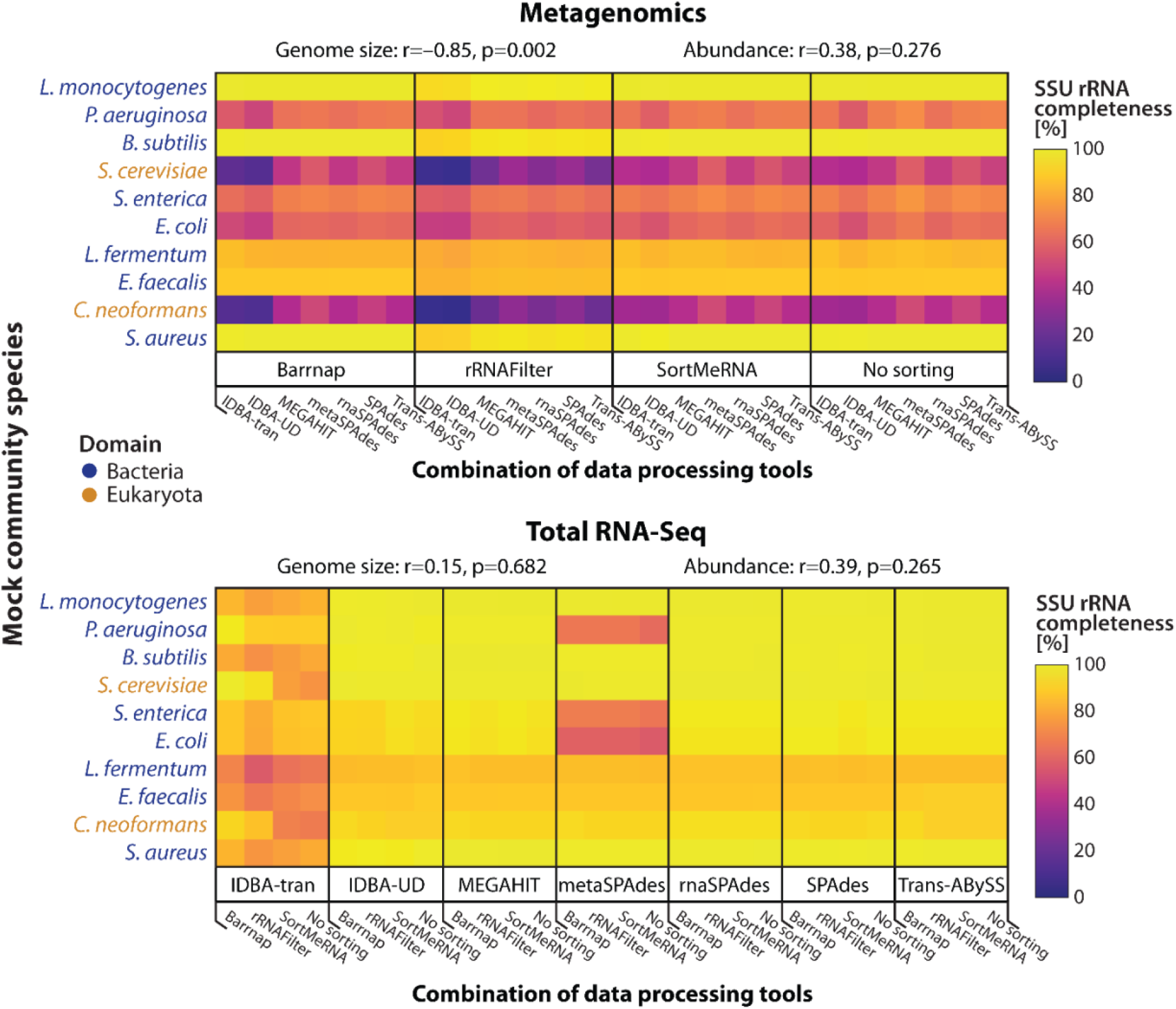
SSU rRNA completeness across mock community species and data-processing combinations. The x-axis of each heatmap represents the combinations of data-processing tools employed, the y-axis represents the ten mock community species, and the colour bar indicates SSU rRNA completeness in percentage. The correlation coefficients and p-value between SSU rRNA completeness and genome size and abundance, respectively, are shown above each heatmap. The combinations are sorted differently for each sequencing method to highlight the tool variations specific to the sequencing methods. Using total RNA-Seq, most species had full-length sequences recovered (indicated by bright yellow) using at least one combination of tools.

The genome completeness of all species was close to 0% for either sequencing method across all utilized data-processing tools and species except for *Listeria monocytogenes*, whose genome was almost 100% complete when applying metagenomics and no rRNA sorting and partially complete when applying metagenomics and rRNAFilter for rRNA sorting (Figure S1).

When testing for significant correlations between each tool and SSU rRNA completeness, the results differed considerably between metagenomics and total RNA-Seq (Figure 5). Each column denotes the correlation between the respective data-processing tool and SSU rRNA completeness, while the rows denote respective species. For metagenomics, the rRNA sorting tool rRNAFilter was significantly and strongly negatively correlated to SSU rRNA completeness across all but one species (p ≤ 0.05; Figure 5, top). Otherwise, no rRNA sorting tool was significantly correlated to SSU rRNA completeness across all species except *L. fermentum*, whose SSU rRNA completeness was significantly positively correlated to no rRNA sorting. Almost no assembly tool significantly correlated with SSU rRNA completeness, except IDBA-UD, which negatively correlated with SSU rRNA completeness in half of the species. For total RNA-Seq, no rRNA sorting tool was significantly correlated with SSU rRNA completeness (p > 0.05; Figure 5, bottom). Only the assemblers metaSPAdes and IDBA-tran were significantly and strongly negatively correlated to SSU rRNA completeness, with IDBA-tran being negatively correlated for three species and metaSPAdes for the other seven.

**Figure 5:**
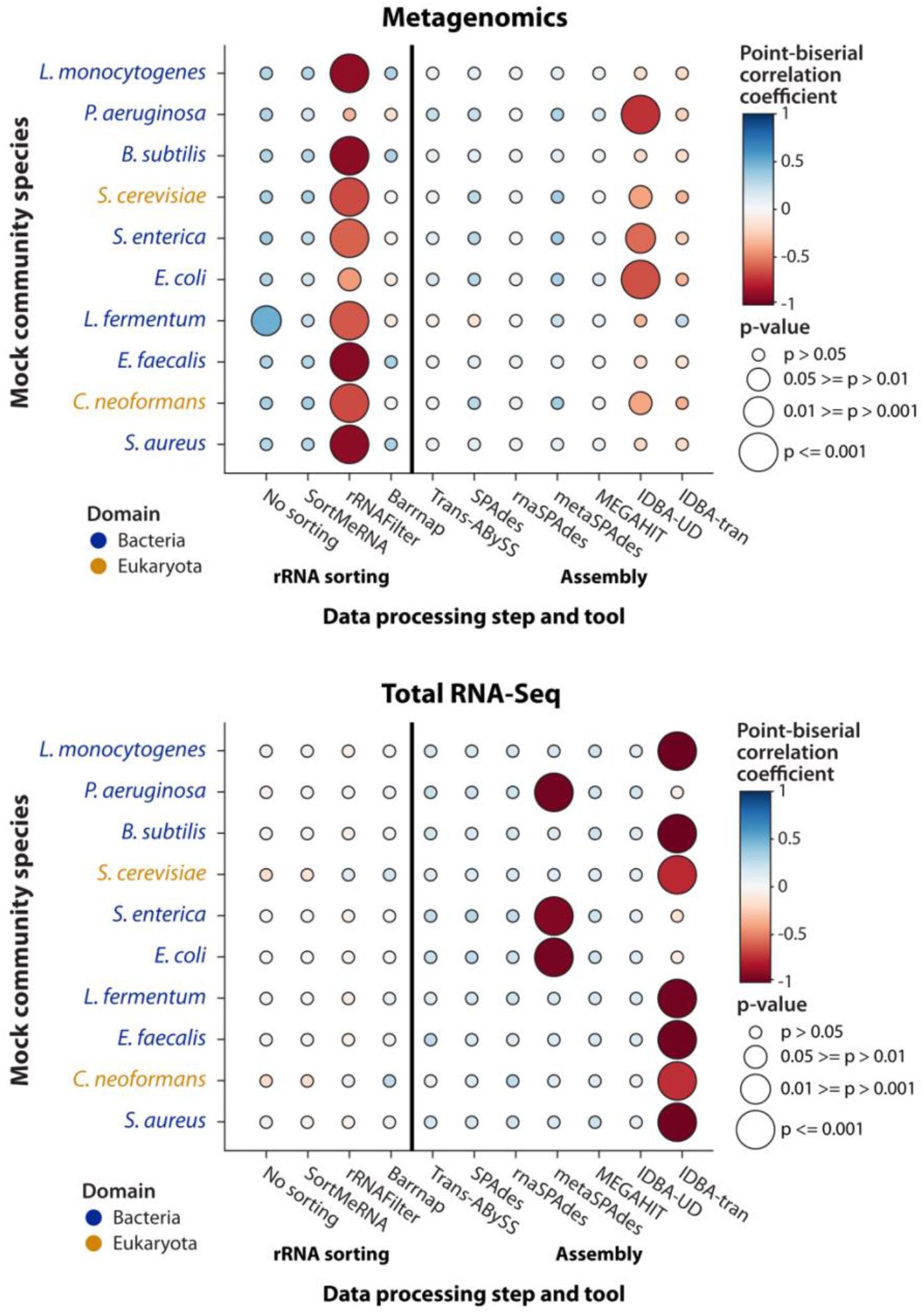
Impact of individual data-processing tools on SSU rRNA completeness. The x-axis of each bubble plot represents the tools employed, the y-axis represents the ten mock community species, the colour bar indicates the point-biserial correlation coefficient between the SSU rRNA completeness and the tools, and the bubble size indicates the significance of the correlations. Large-sized, red bubbles indicate tools that are significantly and negatively correlated with SSU rRNA completeness.

## 4 Discussion

Our results show that total RNA-Seq generates SSU rRNA sequences with completeness equal to or higher than that of metagenomics. This was particularly the case for the two eukaryotic species with genomes much larger than those of the other mock community species, but also for bacterial species with large genomes. In fact, statistical tests confirmed that metagenomics SSU rRNA completeness was significantly negatively correlated with genome size, while this was not the case for total RNA-Seq. This shows that the broad genomic coverage of metagenomics leads to lower coverage of specific genes of interest, such as the SSU rRNA gene, especially with increasing genome size. On the other hand, the SSU rRNA coverage of total RNA-Seq is independent of genome size, confirming that total RNA-Seq naturally enriches the data set for rRNA (Bang-Andreasen et al., 2020; Geisen et al., 2015; Urich et al., 2008).

Surprisingly, the SSU rRNA completeness for three out of the four species with the lowest abundance (relative abundance = 0.015–0.0001%) was on par for total RNA-Seq and metagenomics. In particular, the SSU rRNA of the least abundant species was near-complete for either approach. Statistical tests showed that overall SSU rRNA completeness of neither metagenomics nor total RNA-Seq significantly correlated with species abundance. Given that three out of the four species with the lowest abundance were the only species without near-complete SSU rRNA sequences in total RNA-Seq, it might be possible that the inclusion of more low-abundant species would have revealed a statistically significant pattern, which should be further investigated. Against our expectation that SSU rRNA completeness of total RNA-Seq data is higher for low-abundant species, metagenomics recovered a sufficient amount of SSU rRNA fragments for SSU rRNA reconstruction of low-abundant species. An explanation might lie in the low complexity of the sample. The mock community used in our study contains only ten species. Any difference between metagenomics and total RNA-Seq in terms of SSU rRNA completeness of low-abundant species might become only more prevalent in more complex communities.

The lack of a correlation between abundance and SSU rRNA completeness was in stark contrast to the observed correlation between abundance and genome completeness for metagenomics since the genome of the most abundant species was near-complete while the completeness of all other species’ genomes was close to 0%. This illustrates that the abundance of each species in a community has a significant impact on its genome coverage when applying metagenomics, which is in agreement with other studies showing that successful genome reconstruction of low-abundance species typically requires high sequencing depth (Jin et al., 2022; Merrill et al., 2022). In contrast to that, even the genome completeness of the most abundant species was close to 0% when total RNA-Seq was applied, which further confirms that total RNA-Seq reads are naturally enriched for rRNA and consequently do not provide broad genomic coverage.

SSU rRNA completeness was independent of the applied PHRED score cut-off during quality filtering and trimming. It should be noted, however, that the quality of our sequences was almost exclusively over PHRED 30 according to mean and per sequence quality scores (Fig. S2), so trimming and quality filtering might not have had a great effect in our study but could still have significant effects in studies where only lower-quality data are available.

The methods applied for rRNA sorting impacted the SSU rRNA completeness of metagenomics but mostly not that of total RNA-Seq. In particular, rRNA filter, which sorts sequences based on k-mer abundances, was strongly negatively correlated with metagenomics SSU rRNA completeness, while barrnap, SortMeRNA, and no sorting showed no correlation for any but one species, for which no sorting significantly improved completeness. This shows that extracting rRNA sequences from metagenomics sequences for SSU rRNA reconstruction has no effect on or even decreases completeness and should therefore be omitted, also to save time and computational resources. rRNA sorting had also no effect on total RNA-Seq SSU rRNA completeness, likely because the sequences are already naturally enriched for rRNA, except for the two eukaryotic species, for which rRNA sorting did impact completeness, though only significantly for one species. However, since no specific sorting tool was significantly correlated to SSU rRNA completeness for the two eukaryotic species, we conclude that rRNA sorting can also be omitted for total-RNA-Seq-based SSU rRNA reconstruction, although further research on the impact of rRNA sorting on prokaryotic vs. eukaryotic SSU rRNA reconstruction success is required.

In terms of genome completeness, when metagenomics was applied, only unsorted sequences allowed for complete genome reconstruction of the most abundant species, while genome completeness was close to zero when SortMeRNA or barrnap was applied. This confirms that SortMeRNA and barrnap successfully sort the sequences into rRNA and non-rRNA. Surprisingly, the genome completeness was still comparably high at about 50% on average when applying rRNAFilter, which should not have been the case and indicates that rRNAFilter classified large amounts of non-rRNA sequences as rRNA. Given that rRNAFilter was also strongly negatively correlated with SSU rRNA completeness, we conclude that it should not be used for rRNA sorting in general. Only one other benchmarking study tested rRNAFilter for rRNA sorting and showed that the tool performs poorly in comparison to other rRNA sorting tools (Deng, Münch, Mreches, & McHardy, 2022), which supports our conclusion.

The applied assemblers impacted the SSU rRNA completeness of both metagenomics and total RNA-Seq. No assembler significantly positively correlated with completeness, but for some species, IDBA-UD significantly negatively correlated with metagenomics and metaSPAdes and IDBA-tran with total RNA-Seq. Furthermore, metagenomics-optimized assemblers showed performance similar to that of metatranscriptomics-optimized assemblers for both sequencing methods. These results show that assembly tools mostly do not have a significant impact on SSU rRNA completeness, although specific assemblers can reduce the completeness of some species. Therefore, we suggest using fast and inexpensive established assemblers that have been validated in metagenomics-specific benchmarking studies, specifically MEGAHIT (Awad, Irber, & Brown, 2017; Quince et al., 2017; van der Walt et al., 2017).

## 5 Conclusions

Total RNA-Seq delivered sufficient data for the complete or near-complete reconstruction of all mock community SSU rRNA sequences, and our study confirms that the approach can reconstruct SSU rRNA with equal or better completeness than metagenomics. Furthermore, we show that total-RNA-Seq-based SSU rRNA completeness is independent of genome size, while metagenomics-based SSU rRNA completeness is strongly and significantly correlated with the genome size of taxa in the community. The impact of data-processing tools was overall low, although certain tools resulted in significantly lower SSU rRNA completeness and should be avoided. These results are promising for the high-throughput reconstruction of novel full-length SSU rRNA sequences and the simultaneous application of multiple -omics approaches to routine assessments of ecosystems, which is advocated by multiple studies (Cordier et al., 2021, 2019; Leese et al., 2018; Uyaguari-Diaz et al., 2016). A further comparison of total RNA-Seq with synthetic long-read sequencing, as successfully and effectively applied by Karst et al. (2018), and the short-read-based approach utilized in our study could show which of both approaches is more applicable for routine environmental assessments.

## 6 Acknowledgements

Financial support for this work was provided by the Food from Thought: Agricultural Systems for a Healthy Planet Initiative, by the Canada First Research Excellence Fund (project 000054) through the University of Guelph; grants in Bioinformatics and Computational Biology from the Government of Canada through Genome Canada and Ontario Genomics and from the Ontario Ministry of Economic Development, Job Creation and Trade (project 15401); and from the Natural Sciences and Engineering Research Council of Canada (NSERC Discovery Grant RGPIN-2022-04569).

