## Supplementary material for "Reconstruction of Small Subunit Ribosomal RNA from High-Throughput Sequencing Data: A Comparative Study of Metagenomics and Total RNA Sequencing"

### Supporting information

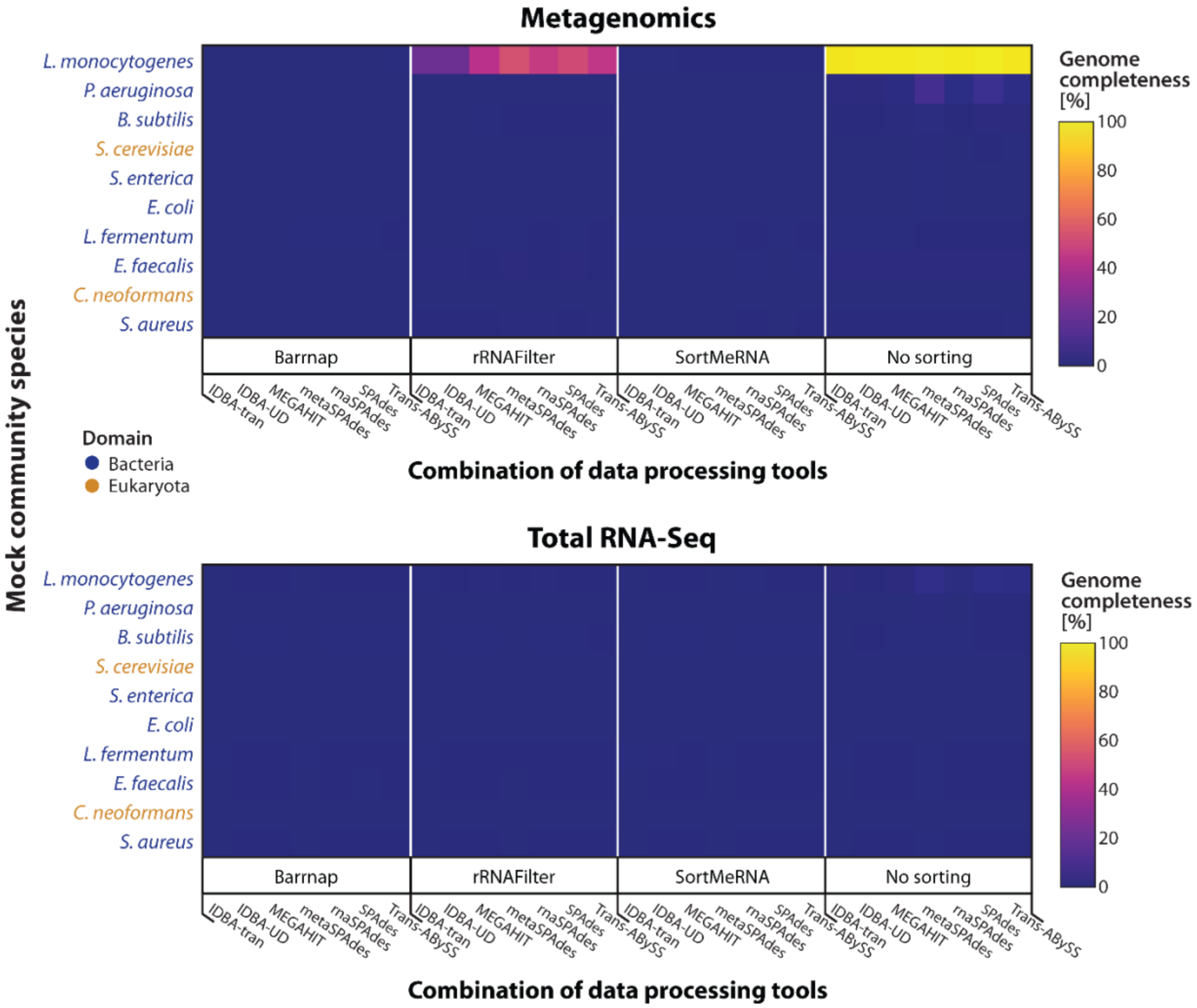


*Supplementary Figure 1: Genome completeness across mock community species and data-processing combinations. The x-axis of each heatmap represents the combinations of data-processing tools employed, the y-axis represents the ten mock community species, and the colour bar indicates SSU rRNA completeness in percentage. Both sequencing methods yielded genome completeness values near zero, with the exception of one abundant species using metagenomics.*

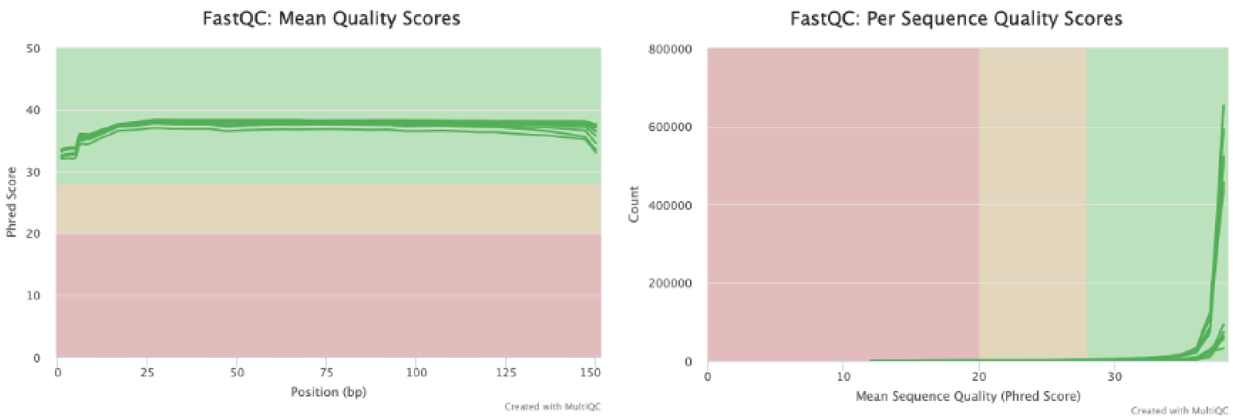


*Supplementary Figure 2: Mean and per-sequence quality scores for metagenomics and total RNA-Seq data. Each line represents forward or reverse reads of one sample (12 in total).*
